# Target interception in virtual reality is better for natural versus unnatural trajectory shapes and orientations

**DOI:** 10.1101/2024.08.16.607980

**Authors:** Sofia Varon, Karsten Babin, Miriam Spering, Jody C. Culham

## Abstract

Human performance in perceptual and visuomotor tasks is enhanced when stimulus motion follows the laws of gravitational physics, including acceleration consistent with Earth’s gravity, *g*. Here we used a manual interception task in virtual reality to investigate the effects of trajectory shape and orientation on interception timing and accuracy. Participants punched to intercept a ball moving along one of four trajectories that varied in shape (parabola or tent) and orientation (upright or inverted). We also varied the location of visual fixation such that trajectories fell entirely within the lower or upper visual field. Reaction times were faster for more natural shapes and orientations, regardless of visual field. Overall accuracy was poorer and movement time was longer for the inverted tent condition than the other three conditions, perhaps because it was imperfectly reminiscent of a bouncing ball. A detailed analysis of spatial errors revealed that interception endpoints were more likely to fall along the path of the final trajectory in upright vs. inverted conditions, suggesting stronger expectations regarding the final trajectory direction for these conditions. Taken together, these results suggest that the naturalness of the shape and orientation of a trajectory contributes to performance in a virtual interception task.

## Introduction

Motion processing is often studied with psychophysics using simple stimuli (such as single dots or random dot kinematograms), canonical 2D motion patterns (such as linear motion, rotation, and optic flow), and predominantly perceptual tasks (Nakayama, 1985; Nishida et al., 2018). Moreover, studies of the neural correlates of motion processing focus largely on the earliest stages of vision (Andersen, 1997; Culham et al., 2001). However, in the real world, visual motion can be more complex and is often used to fulfill ecological functions through integration with other neural systems, including those for visually guided actions (e.g., Land & Tatler, 2009). As cognitive neuroscience research moves to incorporate more naturalistic stimuli and tasks (e.g., Gibson, 1997; Snow & Culham, 2021), more ecological paradigms will enable a better understanding of visual motion processing in everyday life.

One example of an ecological motion scenario is intercepting a projectile such as a thrown ball. In this case, the motion follows the principles of physics, travelling along a parabolic trajectory with an acceleration consistent with Earth’s gravity, *g* (9.81 m/s^2^). Growing evidence suggests that the brain has an internal representation of gravity that influences the processing of moving objects in the natural environment (Delle Monache et al., 2021; Jörges & López-Moliner, 2017; Lacquaniti et al., 2015; Zago et al., 2008). This internal representation (or prior) is formed based on experience with naturally moving objects. In principle, the visual motion system could simply use the target’s initial motion, particularly velocity and acceleration, to predict the target’s path over space and time (e.g., Saxberg, 1987). However, internalization of physics priors would enable better perceptual sensitivity and faster, more accurate actions (Jörges & López-Moliner, 2017; Tresilian, 2005), even in the face of neural processing delays (Zago et al., 2008). Furthermore, there is evidence that people are poor at distinguishing changes in visual acceleration (Gottsdanker et al., 1961; Kreyenmeier et al., 2022; Mueller et al., 2017; Watamaniuk & Heinen, 2003; Werkhoven et al., 1992), suggesting an additional benefit to internalizing physical principles.

Gravitational priors have been found to affect many aspects of behavior including visual perception, gaze tracking, and manual interception (Delle Monache et al., 2015; Diaz et al., 2013; Zago et al., 2009). Notably, cognitive reasoning about physics principles is frequently erroneous (e.g., Hecht & Bertamini, 2000; McCloskey et al., 1980; Shanon, 1976). This stands in contrast to rapid and accurate eye and hand movements toward moving targets, leading to the proposal of a dissociation between cognitive-perceptual processing and action-related processing of moving targets (Tresilian, 1995; Zago & Lacquaniti, 2005). Such a dissociation is consistent with other models of a perception-action distinction in behavior and the underlying neural substrates (Goodale & Milner, 1992; Spering & Carrasco, 2015; Spering & Montagnini, 2011).

Across many studies, performance in perception and action tasks has been shown to be better for motions consistent with downward gravity. Performance is consistently better for falling targets (accelerating at *g*) than rising targets (accelerating at -*g*; Jörges & López-Moliner, 2017; Moscatelli & Lacquaniti, 2011; Senot et al., 2005; Torok et al., 2019). Similarly, performance is better when the trajectory is an upright parabola (with an apex at the top) than an inverted parabola (with a nadir at the bottom; Jörges & López-Moliner, 2019), and when the parabolic trajectory is oriented to the Earth’s vertical compared to when it is rotated up to 90° (Balestrucci et al., 2021). Participants also generally show better performance for targets moving with a natural acceleration (1*g*) compared to zero or other positive accelerations (e.g. 0*g*, ½*g*, 2*g*). This advantage has been found for purely vertical motion (McIntyre et al., 2001; Zago et al., 2004) and projectile motion (Bosco et al., 2012; La Scaleia et al., 2020; Russo et al., 2017), even when the final portion of the object’s trajectory is occluded (Bosco et al., 2012; Delle Monache et al., 2015; Zago et al., 2010), though one study found no clear benefit of 1*g* compared to 0*g* (Jörges et al., 2021).

Despite the emerging consensus that motion direction and acceleration affect performance, there are some possible confounds. Comparisons of accelerations are typically confounded by stimulus durations in that faster accelerations require quicker responses. Another potential confound is the visual field location. With visual fixation or gaze tracking of the target, the region where the target is heading may fall within the lower visual field (LVF) or the upper visual field (UVF). Hand actions are faster and more accurate in the LVF than in the UVF (Brown et al., 2005; Danckert & Goodale, 2001) and brain regions implicated in hand actions show enhancements in the LVF compared to the UVF (Galletti et al., 2003; Rossit et al., 2013), consistent with vision of the hand predominantly in the LVF (Mineiro & Buckingham, 2023).

Natural gravity affects not only acceleration but also the shape of a projectile’s trajectory. Other than inversions of parabolic trajectories (Jörges & López-Moliner, 2019), shape has not been as well studied as acceleration; however, shape distortions could provide an approach to studying gravitational priors without confounds of motion duration and visual field. Participants, even ball players and physicists, are surprisingly poor at cognitive tests of their understanding of projectile shapes (Hecht & Bertamini, 2000; Kozhevnikov & Hegarty, 2001). For example, participants judge circular and sinusoidal curves as being as natural as parabolas (Hecht & Bertamini, 2000). Interestingly, as scene realism improved to include pictorial depth cues, participants judged the veridical trajectories as the most natural (Hecht & Bertamini, 2000). One line of research displayed parabolic trajectories in which the ascent was consistent with *g,* but the descent had accelerations of *g* (normal parabola), 2*g* (generating a narrower parabola with a tighter descent), or 0*g* (generating a wider, more linear descent; Bosco et al., 2012; Delle Monache et al., 2015, 2019). Although effects of acceleration were found, this paradigm changes not just the shape of the trajectory but the duration of the movement. Responses to parabolic trajectories have also been studied in the context of the “outfielder problem”, in which a baseball player must catch a fly ball. However, this particular situation enables strategies that are not necessarily useful for all motion trajectories (Chapman, 1968; Fink et al., 2009; McBeath et al., 1995). Some studies suggest that participants have an expectation of natural trajectories, as people generally perform worse when intercepting objects that are rolling down an unfamiliarly-shaped ramp (Mijatović et al., 2014); they can also predict the trajectory a bouncing object will take based on its elasticity (Diaz et al., 2013). However, how the naturalness of a trajectory affects interception of a moving projectile is unknown.

In this study, we investigated whether trajectories consistent with gravity evoked faster and more accurate manual interceptions than other trajectories even when the duration and visual field of the trajectory were controlled. We were particularly interested in studying manual interception – rapid reaches toward a projectile – as a stepping stone toward further behavioral and neuroimaging studies of reaching under real-world conditions. The project specifically addresses the contribution of trajectory shape, comparing a parabola vs. a control path, and trajectory orientation, comparing upright vs. inverted paths.

We designed a manual interception task in virtual reality (VR). With the advent of high-quality consumer-grade VR, more studies are using VR to study the effects of gravity on perception and particularly action (Fink et al., 2009; Jörges et al., 2021). VR provides compelling immersion and engagement (Adolf et al., 2019; Baurès et al., 2009). Moreover, scene geometry veridically matches the egocentric perspective of an agent in a natural environment. This stands in contrast to some prior studies of gravitational influences on perception, which utilized pictorial cues (Delle Monache et al., 2015; La Scaleia et al., 2020; Moscatelli & Lacquaniti, 2011; Torok et al., 2019). For example, in one study, participants used their gaze to follow a baseball moving through a parabolic trajectory with *g* appropriate to the pictorial geometry rather than real-world physical geometry (Delle Monache et al., 2015). While pictorial physics appear to effectively evoke adequate predictions from gravitational priors for perception, it is possible that systems supporting hand actions may be more tuned to egocentric physics.

We systematically manipulated two features that determine the naturalness of a trajectory: *Shape* (parabola or tent) and *Orientation* (upright or inverted; Figure 1a-d). Participants used their right hand to punch to intercept a virtual projectile. Only one of the trajectories used in our experiment followed an upright parabolic curve (Figure 1a), which adheres to natural laws of gravity; the remaining three trajectories (Figure 1b-d) were less natural. These unnatural trajectories include a tent (Figure 1b), and an inverted condition of both the parabola (Figure 1c) and the tent (Figure 1d). The tent was selected as the unnatural trajectory shape, as its path is symmetrical and could be designed to have the same motion duration as a parabolic trajectory. In this way, the effects of shape could be determined while other features defining physical motion remained constant. Although the inverted tent trajectory is reminiscent of a bouncing ball, only the first (left) half was consistent with the physics of bouncing balls. Under all conditions, the virtual objects moved along a symmetrical path with an acceleration of *g,* such that only trajectory shape and orientation varied between conditions. All motion trajectories took the same amount of time, avoiding the duration confounds present if acceleration is manipulated. To ensure that differences between conditions were not confounded by processing differences between the LVF and UVF, we had participants maintain visual fixation in two locations (Figure 1a-d), so as to place the trajectory in either the LVF or UVF. Varying shape, orientation, and visual field resulted in a 2 x 2 x 2 factorial design for a total of eight trajectory conditions.

**Figure 1.**
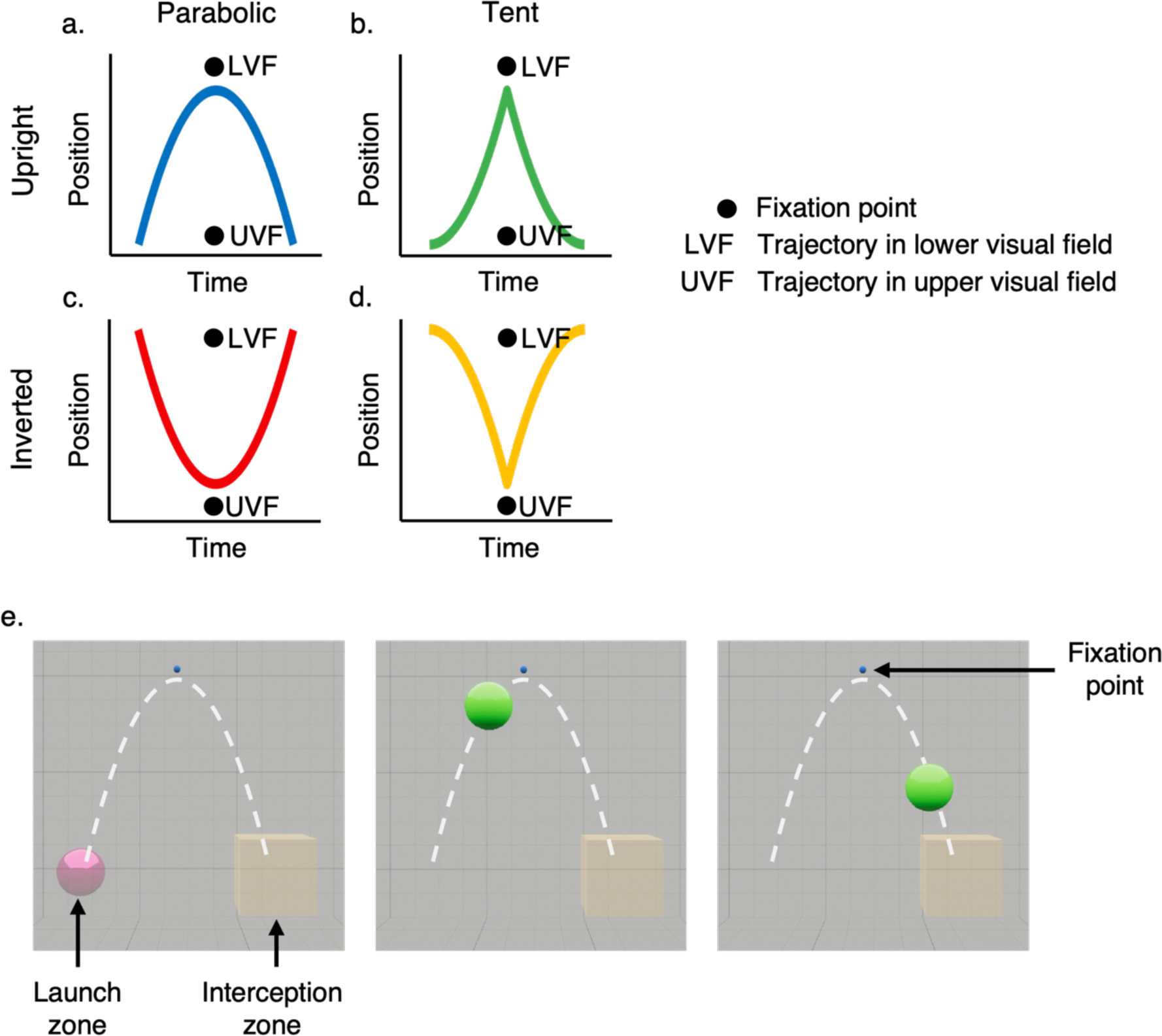
Possible experimental conditions and virtual environment. Participants were tasked with intercepting a virtual ball moving under one of eight conditions. These eight conditions are composed of four possible trajectory shapes which either occur in the lower or upper visual field depending on the location of the fixation point. The fixation point was either at the apex of the trajectory or in line with interception so that interceptive actions were performed in either the upper visual field (UVF) or in the lower visual field (LVF). **(a)** Natural condition of parabolic motion, as predicted by the laws of gravity. **(b)** Unnatural tent-shaped trajectory. **(c-d)** Unnatural, inverted trajectories of both the parabola and tent shapes. **(e)** Virtual environment. An overview of the trajectory for the condition of upright parabola in the LVF is shown. In this condition, the fixation point is placed at the apex of the trajectory. The ball is launched from the launch zone, and participants punch to intercept it once it is in the interception zone. The white dotted line is only for illustration of the trajectory and is not seen by participants.

We predicted that interception would be best – fastest and most accurate – for the most natural trajectory (upright parabola) compared to the unnatural trajectories (inverted and/or non-parabolic), regardless of the visual field in which the trajectory was presented.

## Methods

### Participants

Data were analyzed from 24 participants (12 females and 12 males; ages 18-27), recruited through Western University’s psychology research pool or through advertisements on campus. Data from two additional participants were collected but not used for analysis (based on quality assurance, as described below). All participants were right-handed, as measured by the 10-item Edinburgh Handedness Inventory (Oldfield, 1971). All participants reported normal or corrected-to-normal visual acuity. To ensure that participants had normal depth perception, which could influence target interception, we only included participants with no history of strabismus or amblyopia and with normal stereoacuity (threshold of 40 arcsec or better) as assessed by the Randot Preschool Stereotest^TM^. The study was approved by the Non-Medical Research Ethics Board of the University of Western Ontario and all participants provided informed consent, in accordance with the standards of the Declaration of Helsinki. Participants received either course credit or payment ($15) for their time.

### Overview

As shown in Figure 2a, participants were seated while wearing a virtual reality headset (head-mounted display) that depicted a virtual environment in which a ball could move from left to right along one of four trajectories (Figure 1a-d). Using a controller in their right hand, participants were instructed to “punch” the ball as it passed through an interception zone (as in La Scaleia et al., 2014; Zago et al., 2004). We chose a punching action to prevent participants from adopting a “catching” strategy (moving the hand beneath the ball in anticipation), which would make it difficult to assess spatiotemporal errors. The limited field of view of the head-mounted display (HMD) restricted the maximum height of the trajectory. Due to this restriction and the use of natural physics (acceleration of *g*), the duration of the ball’s movement was short (617 ms), and participants thus had to make rapid movements to intercept it. The rapid nature of these interceptions meant that participants had limited time to use visual cues to correct their movements. As the task was in virtual reality, their hand did not have to stop or slow down as they punched through the ball. To limit the influence of eye movements on interception and therefore the use of different gaze strategies (de Brouwer et al., 2021; Fooken et al., 2021), participants were instructed to maintain fixation throughout the experiment.

**Figure 2.**
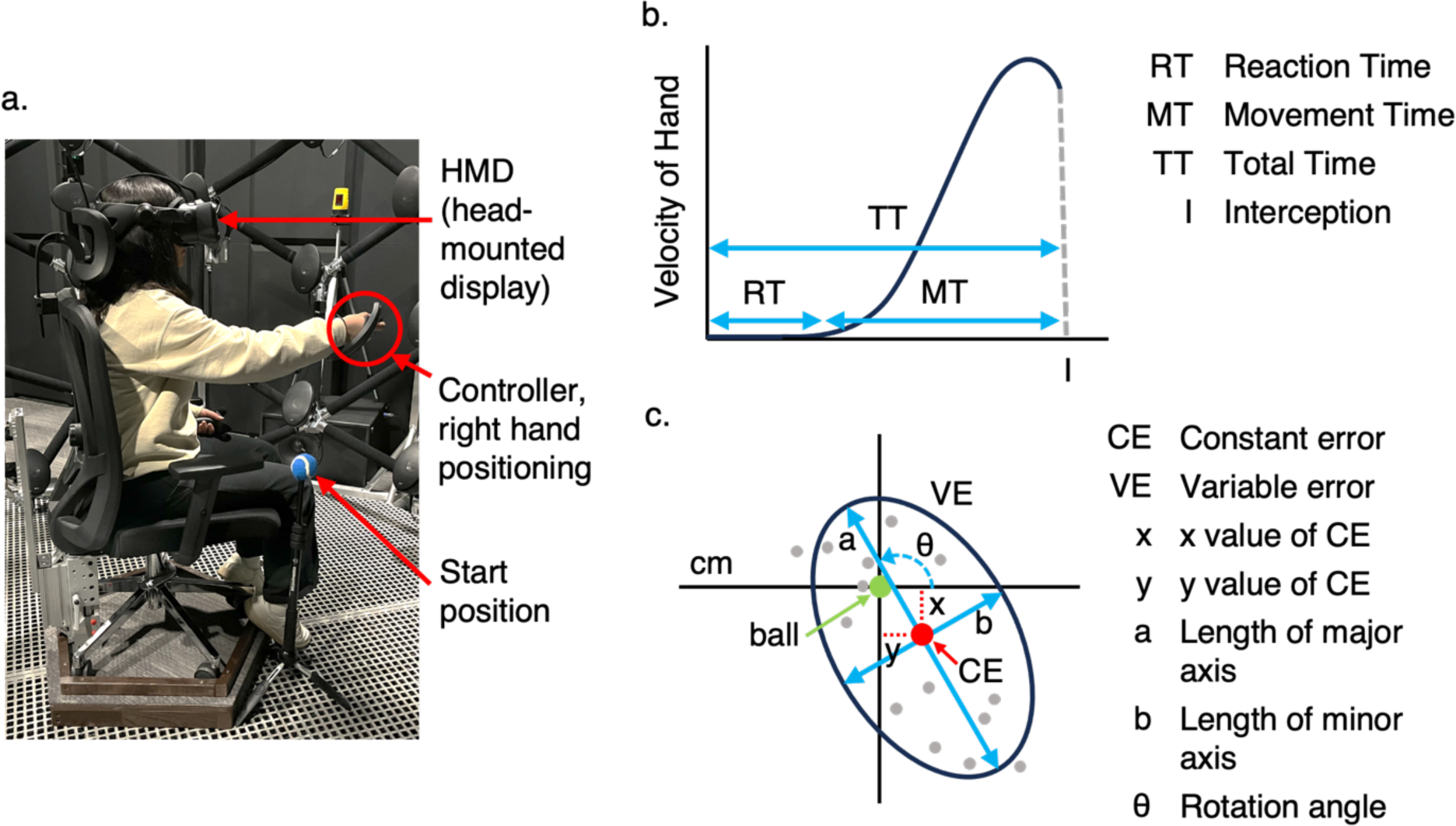
Set up of the testing room and schematic of dependent variables. **(a)** Participant completing the task in the testing room. The participant is wearing the virtual reality HMD and is extending their right arm to intercept the virtual ball with the controller in a punching motion. Next to the participant is a tripod where the participant rests their hand at the start of each trial. **(b)** Schematic of timing variables. Shown is a velocity-time graph that represents the movement of the participant’s hand throughout the trial and depicts how each timing variable is measured. The time of interception is shown as “I.” The shape of the curve is an approximation of the hand-velocity profile during trials. Note that this graph does not represent the position of the ball over time. **(c)** Schematic of spatial error variables, which were computed separately for each condition. The grey circles represent hypothetical reach endpoints from different trials. The constant error (CE), represented as the red circle, indicates the average location of endpoints relative to the ball’s location (represented as the green circle) as the hand passed through the 2D plane of the ball’s trajectory. The x and y components of the constant error are indicated by the dotted red lines. Variable error (VE) was summarized with an ellipse that encompassed 95% of the trialwise reach endpoint locations. The lengths of the major (a) and minor (b) axes and the rotation angle (θ) describe the shape of the variable error ellipse.

We captured measures of timing and accuracy as dependent variables (Figures 2b and 2c). Specifically, timing measures were reaction time (RT) and movement time (MT), which sum to form total time (TT). Examinations of reach velocity plots (Figure 2b) indicated that reach velocity accelerated throughout the movement and peaked around the moment of interception (sometimes just before or just after). This is unlike reaching tasks that involve pointing or grasping, where peak velocity occurs approximately halfway through the movement (Jeannerod, 1984). As such, in our task, measures like time-to-peak-velocity were closely related to MT and are not included here. Measures of accuracy included the percentage of trials in which no errors were made (error trials consisted of trials where either one or various types of errors occurred). Because the ball could be punched at any point during which it passed through the interception zone (time range 517-617 ms for parabolic trajectory, 451-617 ms for tent trajectory), we could not compute simple timing errors. However, as errors in timing would be related to errors in spatial accuracy and precision, we computed constant error (CE) and variable error (VE) for reach endpoints relative to the location of the ball at the time the hand passed through the 2D plane of the ball’s trajectory.

### Details of Task and Setup

During the study, participants wore a Valve Index virtual reality HMD which was adjusted to fit comfortably on each participant. The Valve Index was chosen because it has a relatively large field of view compared to other modern HMDs (Angelov et al., 2020). The Valve Index has a dual-screen display (one display for each eye) with a field of view of 108° (horizontal) × 104° (vertical), a per-screen resolution of 1440 × 1600 pixels, and a refresh rate of 120 Hz. The distance between the screens is adjustable to an interpupillary distance (IPD) range of 58-70 mm. To ensure that the virtual image was focused and that participants could see the image with full binocular vision, participants were instructed to adjust the placement of the HMD’s screens to their IPD, which was measured with the iPhone app *PDCheck AR* (EyeQue Corporation). Participants used a slider on the HMD to adjust the screen positions, which prompted an on-screen pop-up window that actively displayed the current screen distance (in mm) while they moved the slider, allowing participants to know when the screens were at a distance that matched their measured IPD. Participants also held two Index Knuckle Controllers paired to the headset, one in each hand. Positions in space of the HMD and the controllers were monitored through outside-in marker-based tracking, using four external SteamVR 2.0 base stations to track the HMD and controller position at a frequency of 100 Hz. The HMD was calibrated to the space of the testing room for a seated position using the built-in SteamVR calibration process.

Participants sat on a stationary chair (Figure 2a). A headrest was mounted to the back of the chair, and participants were instructed to keep their head against it to maintain a consistent head position throughout the experiment. Between trials, participants rested their right pinkie finger on a tripod with a tennis ball on top, placed to the right of the participant. The height and position of the tripod were adjusted to be comfortable for each participant. Participants positioned their right hand over the controller such that their index finger rested on a rhinestone at the tip of the controller, ensuring consistent hand orientation across participants. The tip of the controller was used to track the position of the participant’s hand in the virtual space. This hand positioning was calibrated and aligned so that the virtual representation of the participant’s hand would match proprioception (as detailed below). Additionally, the most accurate point on the controller is the tip; by positioning the participant’s finger here, we were able to ensure that the finger’s location in virtual space at the time of interception could be accurately monitored.

The experiment software was a custom-made Unity application that implemented calibration, acclimation to the VR environment, and experimental tasks. The Unity editor and game engine were used to develop and render the virtual environment with real-world sizes and distances (whereby 1 Unity unit = 1 m) using Unity’s XR Interaction Tool Kit.

### Experimental Procedure

Participants started the experiment with spatial calibration in the experiment software. During calibration, participants were instructed to manipulate a virtual interception box with their right-hand controller and place it at a reachable distance by extending their right arm at chest height to a comfortable reach position. This distance was used to establish the virtual location of the interception zone, which was centred 5 cm closer than the zone that could be reached comfortably. Both the rest location (atop the tennis ball) and interception zone location were recorded by the application, and a digital sphere was rendered in the virtual location of the rest position to provide participants with a virtual landmark as well. The placement of the rest location and interception zone led to reach trajectories that were upward, forward, and toward the body midline.

After calibration, participants were acclimated to the virtual environment by completing a virtual “fruit-catching” task (somewhat like the video game *Fruit Ninja*, Halfbrick Studios). In this task, fruits were launched in the 3D virtual space at random times. Participants were instructed to use the controller in their right hand to punch these fruits as they appeared in mid-air. The task ended once participants successfully intercepted 30 fruits. This task allowed participants to get familiar with the spatial layout of the scene and their proprioception within the virtual environment. This was done to reduce practice effects during early experimental trials due to being unfamiliar with the controls and with virtual reality environments.

In the experimental task, participants were instructed to be as fast and as accurate as possible when punching with their right-hand controller to intercept a moving ball while it was within the interception box. To launch the ball in each trial, participants pressed a trigger on their left-hand controller. To prevent stereotyped movement onset and ball anticipation, the launch of the ball following the button press was randomly delayed by 500-1500 ms and participants were instructed to not move their hand from the rest position until the ball was visible. Participants engaged in 16 practice trials to get acclimated to the task before completing the experimental trials. During practice trials, participants heard a beep that indicated when the ball was launching, as well as a high-pitched chime if they had successfully intercepted the target. Participants also received visual feedback of how far their hand was from the ball when they attempted to intercept it, represented by a green circle showing where their hand was when they crossed the two-dimensional (2D) plane of the ball’s trajectory and a red circle showing where the ball was at that moment.

Once participants had completed the practice trials, they started the experimental trials. Fifty trials were completed for each of the eight conditions, which amounted to a total of 400 trials in the experiment. During experimental trials, participants did not receive auditory feedback and they were not shown how close their interception was to the target. However, participants were able to see where their hand was in the virtual space (rendered as a small red circle) throughout the whole trial. The moment of interception was defined as the moment at which the participant’s hand crossed the 2D plane of the ball’s trajectory.

### Stimuli

The virtual ball was rendered as a green sphere with a diameter of 10 cm. In all conditions, the ball’s trajectory had a width of 40 cm and an arc height of 40 cm. The ball’s trajectory moved from left to right such that the interception zone always fell in the right hemifield which may have processing advantages compared to the left for right-handed individuals (e.g., Gallivan et al., 2011; Rossit et al., 2013). Trajectories were precomputed in custom code based on the specified height and width and then rendered in Unity. We first computed the trajectories for the upright parabola and inverted tent and then flipped them vertically to generate the inverted parabola and upright tent. The upright parabolic trajectory of the ball followed the laws of physics, with a consistent initial velocity (2.97 m/s) and acceleration of *g* (-9.81 m/s^2^). The inverted tent trajectory was derived by horizontally concatenating two successive parabolic trajectories and showing only the downward phase of the first (left) and the upward phase of the second (right) parabola. This matched many of the low-level features of the two trajectory types, including the time at which the apex/nadir was reached and the total duration. Note, however, that this approach came with two caveats. First, this made the inverted tent trajectory consistent with the trajectory of a perfectly elastic bouncing ball (such that the post-bounce peak was the same as the initial height). As such, it cannot be said to be a truly unnatural trajectory. Nevertheless, the full trajectory of a bouncing ball would normally be seen in the real world, perfectly elastic collisions do not occur in our everyday environments, and most people have less everyday experience with bouncing balls than parabolic trajectories. A second caveat is that the velocity profiles of the two curves differed, with the parabola having high velocities at the start and end and zero velocity at the apex but the tent having the highest velocities just before and after the nadir. As such, the velocity of the ball as it entered and passed through the interception zone would have been higher for the parabola than for the tent. As we will see, however, performance was nevertheless poorer for the tent than for the parabola, which would not be predicted if performance was biased by these speed differences.

The moving ball was shown in a white virtual room that was constructed from six large planes to form a 10 m^3^ cube. The room’s interior walls had a white grid texture applied to help give a sense of position in the space (Delle Monache et al., 2019). The room contained a transparent yellow interception box on the right-hand side (15 cm × 15 cm ×15 cm), a transparent pink spherical launch zone (diameter 10 cm) and a spherical blue fixation point (diameter of 1.5 cm), as shown in Figure 1e. The virtual ball was launched from the pink sphere shown in Figure 1e. Because virtual hands rendered in Unity may not perfectly match the size and positioning of the participant’s real hand, potentially leading to sensory conflicts between vision and proprioception, the participant’s controller was rendered simply as a small red circle.

### Trial Blocking

Because the HMD has a limited field of view, we limited the trajectory heights to ensure that the trajectory would remain in view without necessitating head movements within a trial. When trial blocks switched between visual fields and/or orientation, participants had to move their heads to ensure that they could see the trajectory, the fixation point, and the interception box within their field of view. Thus, the experiment was set up as a 2 (upright/inverted) × 2 (parabola/tent) × 2 (UVF/LVF) factorial design. Trials were blocked by the orientation of motion and the location of the fixation point (in effect blocking by the visual field the trajectory was in). The shape of the ball’s trajectory (parabola or tent) was randomized within both blocks, and block orders were counterbalanced across participants.

Blocking by orientation of motion and visual field resulted in four possible block orders. A given order was repeated twice within a session for a total of eight blocks, such that all trials of one block condition would not occur at the end of the session, when fatigue effects may be more pronounced. Each of the eight blocks was composed of 50 trials, and each participant had a unique trial order within blocks. Although our main interest was in the 2 × 2 × 2 components of the design, to ensure that the various factors were not modulated by practice effects, our initial statistical analysis included a fourth factor distinguishing between the first period (first four blocks) and the second period (last four blocks), leading to a 2 × 2 × 2 × 2 analysis of variance (ANOVA).

### Dependent Variables

To assess how participants responded to natural and unnatural trajectories, we looked at measures of accuracy and speed. Accuracy was calculated as the percentage of trials (out of 50) where no error was made.

We classified several types of errors. In out-of-box errors, the participant made an interception attempt outside of the interception box. Although these trials are counted as errors for the determination of accuracy, they can still be included for other dependent variables. Thus, we counted trials with no errors and out-of-box errors as usable trials. Reach distance too short errors occurred when the participant’s hand did not reach the 2D plane of the interception zone. Overshoot errors occurred when the participant’s hand crossed the plane of the ball’s trajectory first and was then retracted to intercept the ball, meaning that they overshot their reach and were effectively waiting for the ball to land in their hand as opposed to making an accurate interception. Timing errors included reactions that were too fast (< 150 ms), meaning that the hand moved before the ball was visible, or reactions that were too slow (the ball reached the interception zone before the participant moved their hand). Trials with reach distance too short errors, overshoot errors, and timing errors were treated as unusable and discarded from the analysis of timing and spatial error variables.

To assess participants’ movement speed across conditions, we collected the 3D position of the handheld controller over time which allowed for calculation of reaction time, movement time, and total time (Figure 2b). Reaction time was defined as the time between the ball becoming visible and the onset of movement in the participant’s right hand. Movement time was defined as the time between the onset of the participant’s movement and the moment of interception, and total time was defined as the sum of reaction time and movement time. For each participant, median values (across trials) were extracted for the dependent variables of reaction time, movement time, and total time because medians are more resistant to skewed distributions, without requiring arbitrary decisions about outlier removal.

To assess proficiency in intercepting the target, we examined measures of constant error and variable error for each participant for each condition. Constant error was computed by averaging trialwise reach endpoints separately in the x and y directions. Variable-error ellipses were computed for each condition by calculating the covariance of each participant’s reach endpoint locations. The individual covariance matrices were averaged across participants to calculate the group-averaged 95% confidence interval (CI) ellipse. To assess whether the variable error ellipses differed across conditions, we performed statistical tests on the x and y locations of the centres of the ellipses, the angles of orientation of the ellipses (using circular statistics), the lengths of the major axis of the ellipses, and the anisotropy of the ellipses (ratio between the length of the major axis and the length of the perpendicular minor axis; i.e., elongation). Mean values were used for the dependent variables of x and y values of the constant error, and dimensions of the variable error (angle of orientation, major axis length, anisotropy).

For the analysis of timing and spatial error variables, we excluded unusable trials, as described below (See Results: Error Type Distribution).

### Statistical Analyses

Statistical analyses were conducted with the statistical software JASP (version 0.17.1) and Jamovi (version 2.4.7). Data for all analyses can be found at (https://bit.ly/Varon2024Data).

To investigate potential training effects, we initially considered our experiment a 4-factor design with 2 trajectory shapes × 2 trajectory orientations × 2 visual field positions × 2 periods (first period = trials 1-200, second period = trials 201-400). We were most interested in the first three factors; however, given that participants completed 400 trials during the study, training effects (main effects of *Period*) are possible. If training modulated the effects of *Shape*, *Orientation*, or *Visual Field*, we would expect interaction effects with *Period*. Following recommendations that ANOVAs should be corrected for the number of main effects and interactions tested (Cramer et al., 2016), we utilized a Bonferroni-corrected threshold of *p* < .0033 (*p* = .05 / 15 elements = .0033). By this criterion, we found no interactions between *Period* and any of the other three factors (all *p* > .011) for any dependent variable. The ANOVA did reveal a main effect of *Period* on reaction time (*F_1,23_* = 10.8, *p* = .003, *η^2^ _p_* = .32), indicating that participants’ timing improved over the course of the task, but that their improvement did not depend on the trajectory condition.

Because the factor *Period* did not interact with other variables, in the remaining analyses we collapsed data across the two periods and will report the results of a three-factor ANOVA (*Shape*, *Orientation*, and *Visual Field*) for each dependent variable to evaluate the effect of each trajectory condition. We used a Bonferroni correction based on the number of statistical tests within each ANOVA (three main effects, three two-way interactions, and one three-way interaction), yielding a threshold of *p* < .05/7 = .0071 for each dependent variable. Although we could have corrected for the total number of tests across all variables (*p* < .05/63 = .0008), given the already conservative nature of Bonferroni correction, this would inflate the likelihood of Type I statistical errors. We further investigated interaction effects that were found to be significant with *post-hoc* analysis (two-tailed paired-samples *post-hoc t* tests, *p* < .05) to understand the nature of the interaction. *Post-hoc* analyses were used for the interpretation of interactions but were not corrected for multiple comparisons, as the Bonferroni corrections performed at the ANOVA level limited the likelihood of false positives at that stage (Rosenthal & Rosnow, 2008).

### Quality Assurance

Data were screened to ensure that each participant had a sufficiently large number of usable trials. Before analyzing results, we plotted the percentage of usable trials for an initial sample of 24 participants (in JASP statistical software), which flagged one participant as a statistical outlier (with only 13% of their trials being usable compared to a range of 39-93% for the remaining participants). During this initial assessment we also ensured that there were at least six usable trials (12%) for each of the eight experimental conditions. This flagged one participant with no usable trials for one of the eight conditions. The data from both outlier participants were discarded. To preserve counterbalancing, once two additional participants were tested and verified for accuracy (69% and 42% usable trials), their data replaced that of the discarded participants.

The distribution of data for each dependent variable (across participants) was also checked for normality using the Shapiro-Wilk test. Dependent variables that were not normally distributed (where more than two of the eight conditions resulted in *p* < .05) were transformed to achieve normality. A Box-Cox transformation (Osborne, 2010; Sakia, 1992) was applied to accuracy, x-coordinate of constant error, y-coordinate of constant error, length of the major axis of the variable error, anisotropy of the variable error, movement time, and total time. Transformed data were also verified for normality with the Shapiro-Wilk test. For these transformed variables, the Box-Cox transformed data were used for the three-factor ANOVA. The data for these variables were also Box-Cox transformed for the four-factor ANOVA of practice effects, with the same lambda value that was assigned for the three-factor ANOVA. For simpler interpretation, figures will show non-transformed data.

## Results

### Error Type Distribution

We first assessed error types as an indicator of overall task performance, and to later determine how participants’ accuracy (percentage of trials with no errors) was affected by trajectory. Figure 3 shows the frequency of each error type across participants, collapsed across conditions. The most common types of errors were reach endpoints that fell outside the interception zone (“out of box”, 46%), followed by reaches that fell short of the interception plane such that endpoint accuracy could not be determined (“reach distance too short”, 15%), and overshoot errors (12%). Timing errors, in which the hand started to move after the ball entered the interception zone or latency was too short (< 150 ms) were less frequent (5%). Overshoot and timing errors were typically accompanied by out-of-box errors. On average, only 22% of trials did not contain any errors, though there was considerable intersubject variability (range = 1-50%).

**Figure 3.**
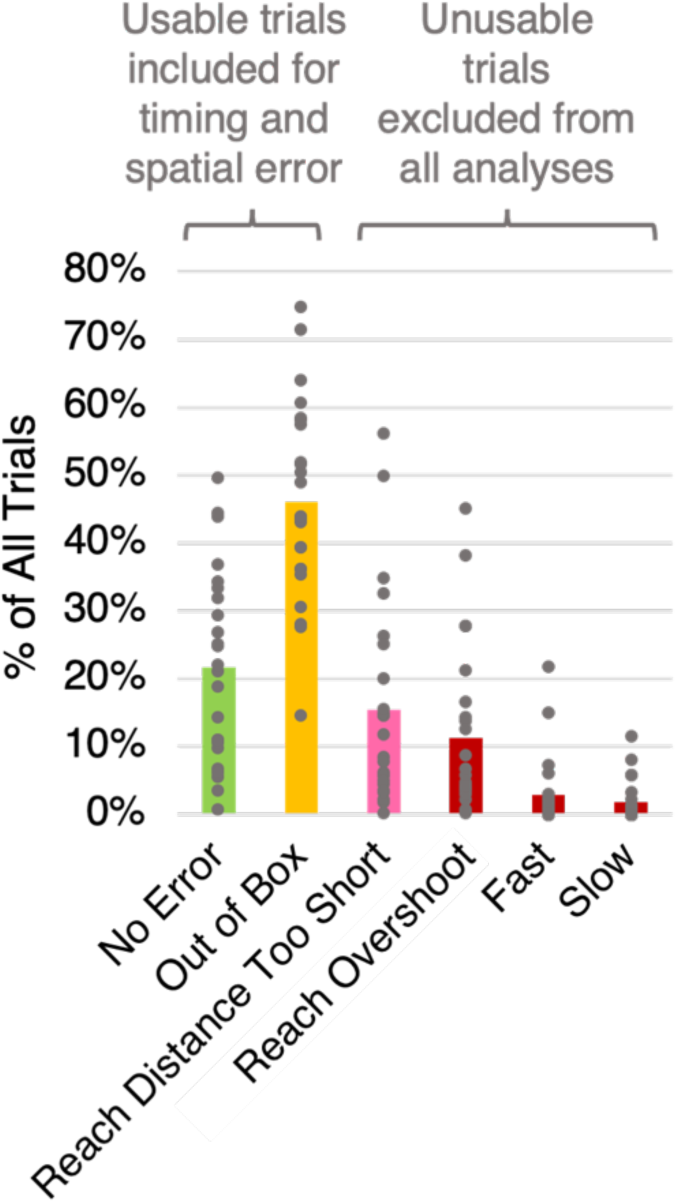
Frequency of each error type across all participants. There were five possible error types, shown in yellow, pink, and dark red. Green represents trials where no errors were committed. The bars represent the frequency of each error type across all trials, collapsed across participants. The distribution of the frequency of each error type across participants is shown by the grey circles. All five error types were counted as errors in the analysis of accuracy, however only errors coloured in pink or dark red (reach distance too short, overshot reach distance, fast, and slow) were considered unusable and were excluded in the analysis of timing variables and spatial error variables. Both green and yellow trials were considered usable trials for the analysis of timing variables and spatial error variables.

For the analysis of accuracy, trials containing any of the five error types were considered error trials. Analyses of all other dependent variables excluded unusable trials (those with too short, overshoot or timing errors), but included out of box trials. On average, we included 68% of trials into these remaining analyses (SD = 15%; range 39-93%).

### Accuracy

We next compared accuracy levels (the percentage of trials with no errors) between conditions. A three-factor ANOVA on the data revealed an interaction between *Shape* and *Orientation* (*F_1,23_* = 54.5, *p* < .001, *η^2^_p_* = .70), an interaction between *Orientation* and *Visual Field* (*F_1,23_* = 10.4, *p* = .004, *η^2^_p_* = .31), and a main effect of *Shape* (*F_1,23_* = 22.1, *p* < .001, *η^2^_p_* = .49) on accuracy. As shown in Figure 4a, the interaction between *Shape* and *Orientation* is explained by poorer accuracy when the trajectory was tent-shaped and inverted (yellow symbols) compared to the other three conditions. One possible explanation is that performance for the inverted tent may be worse because it evokes an impression of a bouncing ball, but the physics of its trajectory is not entirely consistent with that situation in the real world. The absence of a three-way interaction between parabola, tent, and visual field suggests that the effects observed were not simply explained by visual field differences. There was an interaction between *Orientation* and *Visual Field*, such that accuracy was higher in the LVF than UVF, but only for upright trajectories (Supplementary Figure 1), consistent with past reports of a LVF advantage for hand actions (Brown et al., 2005; Danckert & Goodale, 2001), but only for the more natural orientation of motion. However *Visual Field* did not interact with *Shape* or the combination of *Shape* and *Orientation*.

**Figure 4.**
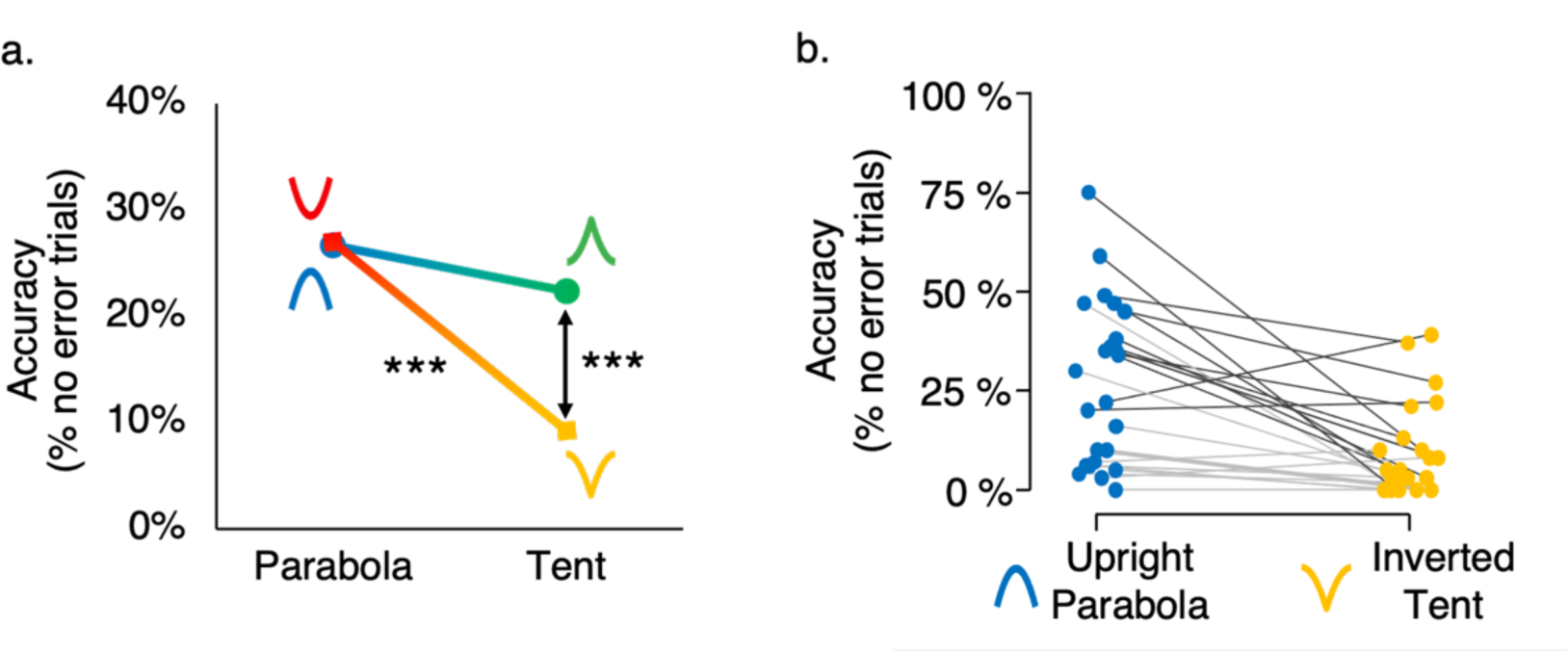
Accuracy across conditions and across participants. * *p* < .05, ** *p* < .01, *** *p* < .001 after *post-hoc* analysis. **(a)** Interaction effect between *Shape* and *Orientation* on accuracy, collapsed across visual field. Participants made more errors when intercepting a ball moving along an inverted trajectory, but only when it followed the tent shape. The schematic beside each data point represents the condition it corresponds to. **(b)** Distribution of accuracy scores across participants between the most natural (UP, upright parabola) and most unnatural (IT, inverted tent) conditions. Each line represents a single participant. Participants were ranked by overall accuracy, and the top 50% of participants are represented by the darker grey lines. Accuracy was largely variable across participants, as some show a strong advantage in natural trials compared to unnatural, while others scored low across all conditions. However, there is a general trend of lower accuracy for inverted tent trials compared to upright parabola trials.

### Individual Differences in Accuracy

Given that there was large individual variability in the percentage of accurate trials across participants (Figure 3), we examined individual differences and their potential impact on the results. Figure 4b shows the percentage of trials with no errors across participants between the most accurate condition (upright parabola) and the least accurate condition (inverted tent), collapsed across visual field. Accuracy across trials was highly variable between participants (range: 0% - 75%). Notably, some participants exhibited very few accurate trials regardless of trajectory shape, while others showed a substantially higher number of accurate trials for the upright parabola shape compared to inverted tent trials. Figure 4b indicates that participants with higher accuracy overall also showed a larger difference between the two conditions (i.e., their lines have a steeper slope), indicating a possible floor effect in performance. However, we observed only a statistical trend in the correlation between overall accuracy and the accuracy difference between upright parabola and inverted tent (*r* = 0.40, *p* = .051).

To ensure that a potential floor effect in performance did not substantially impact our results, we performed additional analyses. We ranked participants by their overall accuracy collapsed across all conditions (as indicated by dark grey and light grey lines for the top and bottom halves, respectively, in Figure 4b). For each dependent variable, we then performed ANOVAs with rank (the 12 participants with the highest accuracy vs. the 12 participants with the lowest accuracy) as a between-subjects variable combined with the other within-participants variables *Shape*, *Orientation* and *Visual Field*. Although split-half analyses have less power than correlational analyses (which do not artificially dichotomize continuous variables; e.g., Altman & Royston, 2006), because of the factorial nature of our design, splitting enabled an assessment about whether main effects and interactions were affected by individual differences in accuracy. The split-half analysis did not identify any significant two-way interactions between rank (high or low accuracy) and any independent variable across any of the dependent variables (all *p* > .006, consistent with a correction for multiple comparisons of *p* > .0033 required for significance of a four-factor ANOVA). Even the pattern of results for the dependent variable of accuracy was not particularly affected by overall accuracy (no interactions between rank and other variables). Indeed, an analysis of the participants whose accuracy was below the median showed qualitatively similar results as the full sample. If anything, our effect sizes for accuracy likely would have been even larger if more participants had performed with high accuracy and the effect size had not been truncated by the floor effect.

### Spatial Errors

We next evaluated spatial errors to determine participants’ proficiency in intercepting the target, and to determine whether there were any patterns in the participants’ reaches. For this and all following analyses, we included both trials with no errors and out-of-box errors. To assess spatial errors, we generated average 95% CI ellipses centred on the average x,y locations of the interception locations for each condition as shown in Figure 5a. Visual inspection of this figure suggests several notable features. First, the constant errors indicate the endpoints were similar across conditions along the horizontal (x) axis, always to the right of the ball, consistent with the left-right direction of the ball’s motion. However, constant errors differed between conditions along the vertical (y) axis, with lower positions for the upright conditions than the inverted conditions. Participants may be accustomed to aiming below a ball in order to intercept it in everyday life; given that the upright parabola condition is more natural than the upright tent, the effect of anticipating the ball’s future location is more pronounced in this condition. Second, the pattern of variable errors indicated differences in the elongation and orientation of the error ellipses. Upright conditions showed trial endpoints scattered along the approximate orientation of the final trajectory. Presumably, in the upright conditions, participants were more attuned to the axis of the ball’s final trajectory, especially when the trajectory was parabolic. During inverted conditions, error patterns were less elongated suggesting less influence of systematic errors. All of these effects were supported by statistical testing as follows.

**Figure 5.**
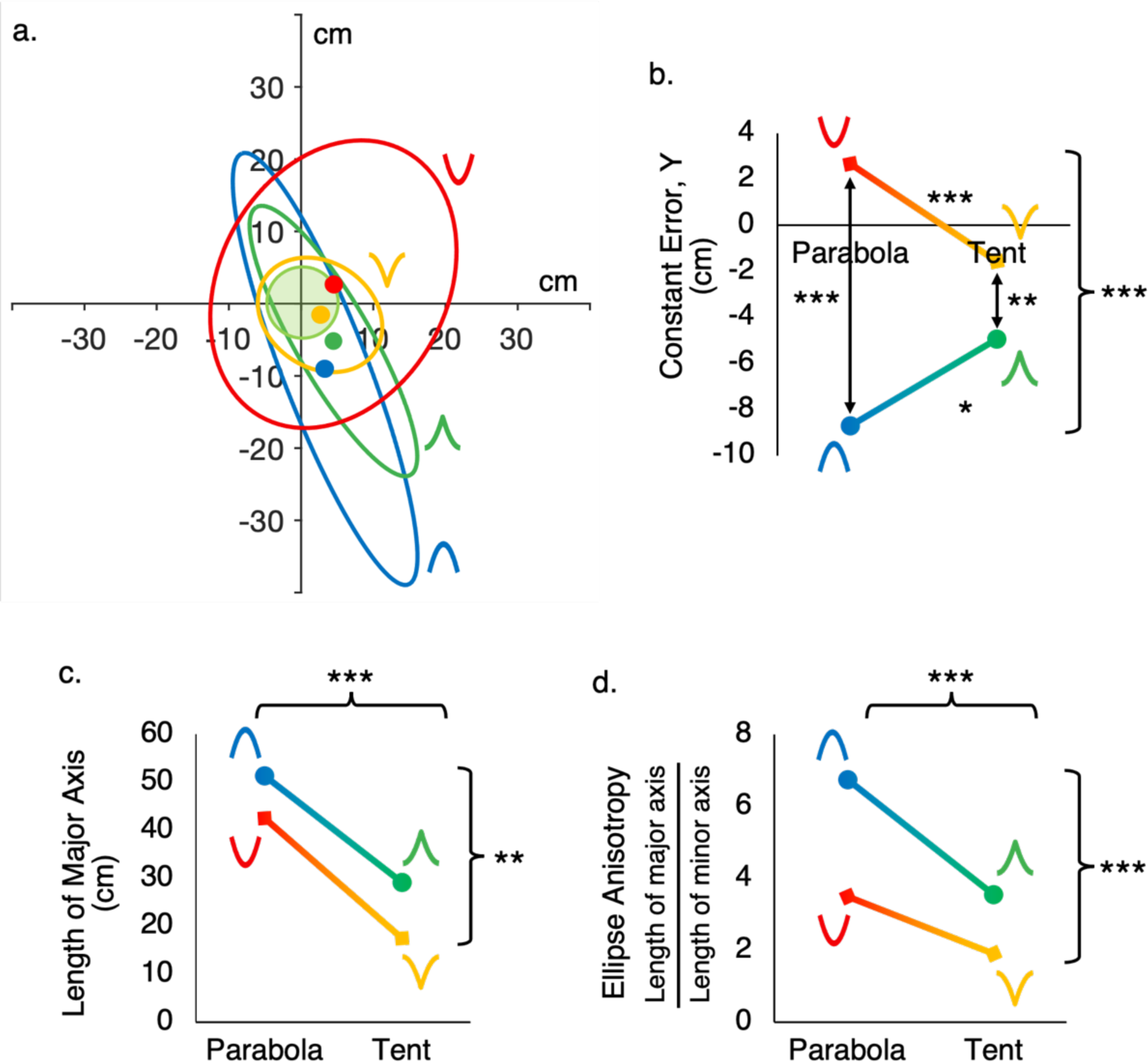
Constant and variable error across conditions. The effect of the different trajectories on spatial error variables, collapsed across visual field, are shown in this figure. The schematic next to each data point depicts the trajectory shape and orientation it corresponds to. Main effects are shown with brackets, and interaction effects are shown with arrows and stars above the lines. * *p* < .05, ** *p* < .01, *** *p* < .001 after *post-hoc* analysis. **(a)** Interception endpoint 95% confidence interval ellipses across conditions. Each ellipse represents the variable error for a given condition centred on its respective constant error, which is represented by the small circles. The colours of the constant error circles and their corresponding variable error ellipses are matched. The constant error was measured relative to the center of the ball at the moment of interception. The transparent green circle at the center, (0,0), is a to-scale representation of the ball, which has a radius of 5 cm. **(b)** Interaction between *Orientation* and *Shape* for constant error along the y direction. **(c)** Length of the major axis of the variable error ellipse. A main effect of *Orientation* and a main effect of *Shape* were identified. **(d)** Anisotropy of the variable error ellipse. A main effect of *Orientation* and a main effect of *Shape* were identified. Anisotropy was computed by dividing the length of the major axis by the length of the minor axis of each ellipse.

A three-factor ANOVA on the data for the average x location of interception identified no significant main effects or interactions; however, the average position across conditions was significantly to the right of the ball’s location (one-tailed *t* test comparing x values vs. 0, *t_23_* = 5.77, *p* < .001, *d* = 1.18). A three-factor ANOVA on the data for the average y location of interception (Figure 5b) identified a three-way interaction between *Orientation*, *Shape*, and *Visual Field* (*F_1,23_* = 14.03, *p =* .001, *η^2^_p_* = .38), along with an interaction between *Orientation* and *Shape* (*F_1,23_* = 19.51, *p <* .001, *η^2^_p_* = .46), an interaction between *Orientation* and *Visual Field* (*F_1,23_* = 30.85, *p <* .001, *η^2^_p_* = .57), and a main effect of *Orientation* (*F_1,23_* = 29.15, *p <* .001, *η^2^_p_* = .56). Inspection of the three-way interaction revealed that the modulation by *Visual Field* did not qualitatively alter the interpretation of the interaction between *Orientation* and *Shape* (Supplementary Figure 2a). The interaction supports the observation that average reach locations were lower for upright than inverted conditions, though the difference was modulated by *Shape*. The two-way interaction between *Orientation* and *Visual Field* is depicted in Supplementary Figure 2b for completeness.

Statistical testing of variable-error ellipse properties supported the observations from Figure 5a. A three-factor ANOVA on the data for the length of the major axis (Figure 5c) identified a main effect of *Orientation*, with longer ellipses for upright than inverted conditions (*F_1,23_* = 8.68, *p =* .007, *η^2^_p_* = .27) and a main effect of *Shape* with longer ellipses for parabola than tent (*F_1,23_* = 99.89, *p <* .001, *η^2^_p_* = .81), but no interactions. Thus, for the more natural conditions, upright and parabolic trajectories, participants likely benefitted from stronger expectations about the ball’s trajectory. A three-factor ANOVA on the data for the anisotropy of the ellipses (i.e., whether they are more elongated or circular; Figure 5d) identified a main effect of *Orientation* (*F_1,23_* = 15.47, *p <* .001, *η^2^_p_* = .40) with higher anisotropy for upright conditions and a main effect of *Shape* (*F_1,23_* = 64.03, *p <* .001, *η^2^_p_* = .74) with higher anisotropy for parabola conditions. The most natural orientation, the upright parabola, yielded the highest anisotropy, providing statistical support for the observations from Figure 5a regarding the major axis.

A two-factor ANOVA (using the Harrison-Kanji test for circular statistics; factors of *Orientation* and *Shape*) on the angle of the ellipse identified an interaction between *Orientation* and *Shape* (*F* = 7.45, *p* = .007), along with a main effect of *Orientation* (*F* = 13.52, *p* < .001) and a main effect of *Shape* (*F* = 22.76, *p* < .001). These results provide statistical support for the visual observations from Figure 5a that endpoints for the upright condition were scattered along the direction of the ball’s trajectory, while during inverted conditions they were scattered in the direction of the ball’s trajectory for parabola but not tent trials.

### Reaction Time, Movement Time, and Total Time

Lastly, we assessed reaction time, movement time, and total time to identify patterns and differences in the timing of participants’ reaches. As total time of a trial is the sum of reaction time and movement time, the results of these three dependent variables will be discussed together. A three-factor ANOVA on reaction time identified a significant main effect of *Shape* (*F_1,23_* = 67.0, *p* < .001, *η^2^_p_* = .19) and a significant main effect of *Orientation* (*F_1,23_* = 63.9, *p* < .001, *η^2^_p_* = .31). No interaction effects were identified. As shown in Figure 6a, unnatural *Shape* and unnatural *Orientation* led to slower median reaction times, although the absence of an interaction suggests that the two factors are additive (but not synergistic). No significant effect was found for *Visual Field*, indicating that the effect of a slower reaction time caused by inverting the trajectory was not a result of the visual field in which participants performed interceptions.

**Figure 6.**
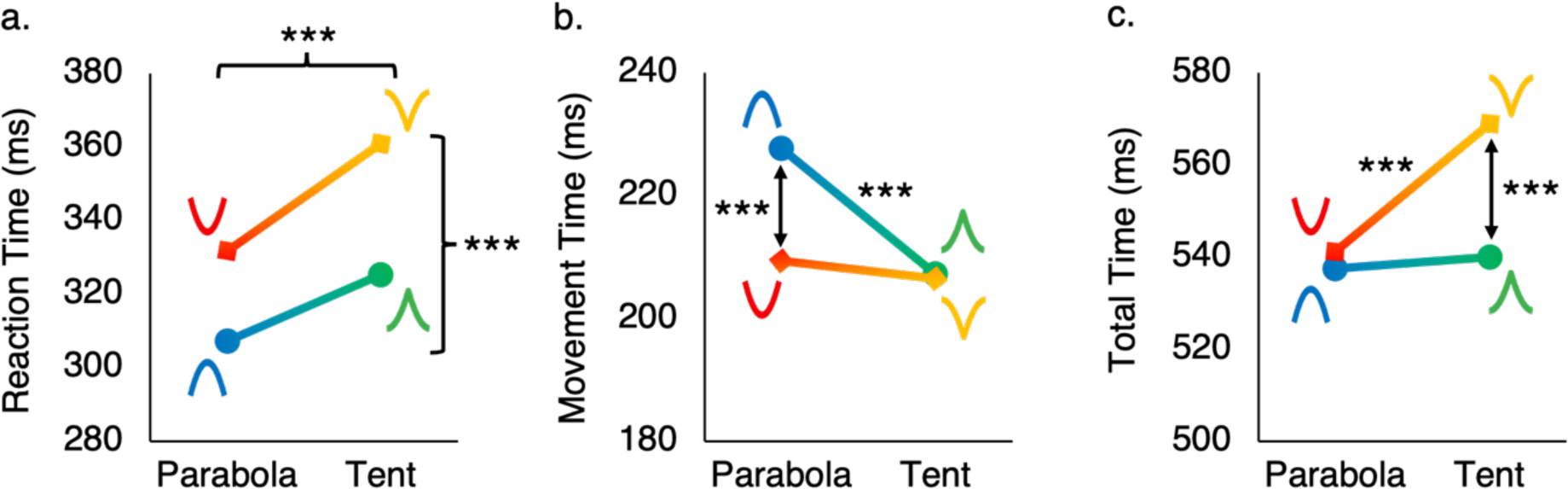
Scores on timing dependent variables across conditions. The effect of the trajectory shapes on timing variables, collapsed across visual field, are shown in this figure. The schematic next to each data point depicts the trajectory shape it corresponds to. Main effects are shown with brackets, and interaction effects are shown with arrows and stars above the lines. * *p* < .05, ** *p* < .01, *** *p* < .001 after *post-hoc* analysis. **(a)** Median reaction time. A main effect of shape and a main effect of orientation were identified. **(b)** Median movement time. An interaction between shape and orientation was identified. **(c)** Median total time. An interaction effect of shape and orientation was identified.

For analysis of median movement time, a three-factor ANOVA on the data identified a significant interaction between *Shape* and *Orientation* (*F_1,23_* = 27.6, *p <* .001, *η^2^_p_* = .55), an interaction between *Orientation* and *Visual Field* (*F_1,23_* = 8.8, *p* = .007, *η^2^_p_* = .28), and a main effect of *Shape* (*F_1,23_* = 9.9, *p* = .005, *η^2^_p_* = .30). As shown in Figure 6b, movement time was significantly slower for the most natural condition (upright parabola) than the other three conditions. Although this finding may initially seem counterintuitive, it may reflect a sensible strategy for interception during unnatural movements. That is, in the face of less intuitive trajectories, participants may take longer to gather information before initiating the movement (reaction time) but then need to perform the movement faster to reach the interception plane in time.

The interaction between orientation and visual field was not a key finding but is depicted in Supplementary Figure 3 for completeness. This interaction revealed that movement time is significantly longer for interceptions in the UVF for upright trajectories compared to the other three conditions.

For analysis of median total time, a three-factor ANOVA on the data identified an interaction between *Shape* and *Orientation* (*F_1,23_* = 24.6, *p* < .001, *η^2^_p_* = .52), as well as significant main effects of both *Shape* (*F_1,23_* = 27.7, *p* < .001, *η^2^_p_* = .55) and *Orientation* (*F_1,23_* = 16.6, *p* < .001, *η^2^_p_* = .42). This relationship is exhibited in Figure 6c, which shows that total time was greater for the inverted tent trial than the three other conditions. This result suggests that while participants may have compensated for a slow reaction time with a fast movement time in order to maintain a consistent total time for two of the unnatural conditions (upright tent and inverted parabola), this compensation was not sufficient to overcome the delay in reaction time imposed by the inverted tent, where both *Shape* and *Orientation* are unnatural.

## Discussion

We found that people are better at intercepting objects that move along natural trajectories compared to unnatural trajectories in terms of both timing and accuracy. The combination of an unusual shape with an unusual orientation in the inverted tent trajectory showed particularly poor performance in terms of both accuracy (Figure 4a) and total timing (Figure 6c), with interactions that could not be explained by orientation and shape alone. Although the total time differed only for the inverted tent compared to the other three conditions (Figure 6c), a breakdown of timing suggested differences in strategy across the four conditions. One possible explanation for the especially poor performance with the inverted tent condition is that it evokes expectations consistent with a bouncing ball that were not perfectly matched by the stimulus. Reaction time (Figure 6a) was fastest for upright conditions compared to inverted trajectories and for parabolas compared to tents. Participants appeared to benefit from the early start on the upright parabola trajectory, which enabled slower reaches (Figure 6b). Even though slower reach movements for the upright parabola might be expected to yield higher spatial accuracy compared to other conditions, this was surprisingly not the case. Rather, a detailed analysis of spatial errors (Figure 5a) showed that for the upright conditions, particularly the parabola, participants’ endpoints were scattered along the axis of the final trajectory and landed lower on average. The breakdown of timing and spatial error variables suggests that participants had strong expectations of how a ball would move for an upright trajectory, particularly a parabolic one, consistent with everyday experience. As such, they could react faster and plan their reaches to fall along the ball’s final trajectory, erring on the side of landing below. Effects of trajectory shape and orientation were generally not modulated by the visual field of presentation.

The results of our study provide evidence that people have stronger expectations for parabolic trajectories compared to other trajectory shapes. The slower reaction time for tent compared to parabola trajectories may reflect a need for additional visual input before generating a movement when faced with unnatural trajectories that deviate from a gravitational prior. Our finding that endpoints were oriented along the ball’s trajectory for the upright parabola condition further suggests that participants had a strong expectation for a parabolic trajectory shape that facilitated extrapolation along the axis of movement. These results corroborate and extend past findings that trajectory shape influences behavior for balls travelling down ramps (Mijatović et al., 2014) and for parabolic trajectories that were distorted by changes in acceleration (*g* vs. other values; Bosco et al., 2012; Delle Monache et al., 2015). Specifically, our results show stronger gravitational priors for parabolic compared to non-parabolic trajectories, even when the trajectory duration is equivalent. We also found that gravitational priors were stronger for upright compared to inverted parabolic trajectories during manual interception (in terms of reaction time and endpoint errors). This finding aligns with previous literature that found faster reaction times for the interception of a target moving downwards compared to upwards (Senot et al., 2005), and better duration discrimination for downward moving targets compared to upwards (Moscatelli & Lacquaniti, 2011; Torok et al., 2019).

We did not find a significant influence of visual field on performance, consistent with smoother pursuit for downward moving targets compared to upwards, regardless of the visual field of the movement (Ke et al., 2013). While these prior studies utilized purely vertical motion, only one study examined the effect of orientation for parabolic trajectories, finding improved smooth pursuit and a faster reaction time for upright compared to inverted parabolas (Jörges & López-Moliner, 2019). Our results suggest upright parabolic trajectories also enhance expectations during manual interception.

We were surprised to find particularly poor accuracy and interception time for the inverted tent trajectory. Although we had not explicitly planned this in designing the trajectories, the inverted tent trajectory appears *similar* but not identical to that of a bouncing ball. Though similar, the tent trajectory is not a perfect match because the post-bounce height would only match the pre-bounce height, which it did in our study, for a perfectly elastic ball. Additionally, our virtual ball did not bounce off any visible floor before making its ascent. It is also unclear how much experience typical participants have with bounce trajectories compared to parabolic ones. Thus, it may be that the similarity to a bouncing ball invoked weak or incorrect expectations about its path that impaired performance. These intriguing results pose new questions about how expectations about gravity, object properties like elasticity, and experience with bouncing balls from sports affect interception of bouncing or rebounding objects in VR, which have been studied with gaze (Diaz et al., 2013; Land & McLeod, 2000; Shalom et al., 2011), perceptual tasks (Warren et al., 1987), simulated interception (Marinovic et al., 2012), and hitting bouncing balls in a cricket task (Müller & Abernethy, 2006).

Our approach highlights that gravitational priors can also be revealed by extending the analysis of errors from simple measures like absolute error (Euclidean distance between endpoint and target) to more complex measures that quantify constant and variable error in at least two dimensions (see also, Russo et al., 2017). Most notably, the upright parabola, the condition with the strongest prior, yielded endpoints that fell along a narrow ellipse aligned along the axis of the final trajectory. Moreover, average endpoints (constant error) were lower for upright than inverted trajectories, consistent with a strategy of motion extrapolation.

### Limitations

A potential concern with our study is that accuracy on the task was low, with only 22% of trials on average containing no errors. However, a substantial number of trials – 46% on average – were classified as errors only because the endpoint fell outside the designated interception zone (“out of box” errors). Moreover, spatial error analysis revealed that the errors committed were systematic, falling along the final trajectory axis, especially in the most natural upright parabola condition, suggesting that participants’ reaches were aligned to the ball’s trajectory. Thus, while these trials were inaccurate in that participants did not aim within the interception zone, the pattern of responses nevertheless revealed expectations about the ball’s trajectory.

Accuracy was likely affected by the limitations of the VR HMD field of view (104° vertically), unlike in the real world where the eyes have a large field of view and head movements are less cumbersome. As HMDs with larger fields of view become available, this would enable higher parabolas (over longer trials) and more natural gaze including not just eye but head movements.

Another limitation is that the acceleration profiles differed between parabola and tent trajectories as a result of how the trajectories were constructed. Specifically, while the parabola begins and ends with a high velocity, reaching zero velocity at the midpoint, the tent begins and ends with zero velocity and reaches its highest velocity at the midpoint. As such, the ball spent more time in the interception zone during tent trials compared to parabola trials. Although this confound would lead to the prediction of better performance for the tent than the parabola trajectories, in fact, the inverted tent condition yielded the worst performance in terms of accuracy, reaction time, total time, and orientation of the variable error ellipse. In contrast, the upright parabola yielded the best performance for reaction time and the shape of the variable error ellipse. Thus, our results would likely be even stronger if the confound were eliminated.

### Future Directions

We developed a novel VR paradigm for manual interception that could easily be extended to build upon the discoveries here. Our paradigm allows participants to make interceptive actions (as in reality) that are likely more effective at tapping into natural behaviors than more simulated tasks such as intercepting a moving target on a computer screen using a mouse and cursor (Zago et al., 2004). Furthermore, VR still enables examinations of intuitive physics through the violation of the laws of physics (which is not possible in reality on Earth).

The most interesting discovery is that visuomotor expectations are enhanced by trajectory shapes consistent with gravity. Future experiments could examine this phenomenon further, including trajectories that are better matched for low-level properties and trajectories that are considered cognitively plausible by participants (Hecht & Bertamini, 2000). This phenomenon could also be extended by studying its role in bouncing motion, and evaluating to what extent a natural shape matters in the perception of bouncing. With newer HMDs incorporating eye tracking, VR would also offer the possibility to investigate whether gravitational expectations are similar for gaze tracking, manual interception, and perceptual-cognitive tasks. Our paradigm could also be used to investigate the contributions of other factors, including the contribution of pictorial context (Miller et al., 2008; Moscatelli & Lacquaniti, 2011). Given our findings of high interindividual variability, it would also be interesting to examine the effects of expertise, such as experience with video games, juggling, or sports, in modulating performance and expectations based on the natural world.

Given that visual neuroscience has traditionally viewed motion and shape perception as being processed in distinct brain regions and networks, another interesting question is how naturalistic motion trajectories are processed in the brain. Past studies have found effects of upward vs. downward motion (Indovina et al., 2005; Miller et al., 2008) and pictorial cues to gravitational acceleration (Miller et al., 2008) in brain regions associated with motion perception (MT+/V5) and vestibular processing. Moreover, target interception for natural (1*g*) vs. unnatural (-1*g*) was perturbed using transcranial magnetic stimulation to the temporoparietal junction (Bosco et al., 2008). Although numerous studies have investigated shape defined by motion (e.g., Handa & Mikami, 2018), only a few studies have investigated “the shape of motion” – that is the trajectory – and its naturalism. Neuroimaging research could be extended to compare natural vs. unnatural trajectory shapes and to include interception tasks and not just perceptual ones. However, given the necessity with fMRI to have participants lay supine, which would distort the vestibular representation of gravity, and remain relatively still in a confined space, other techniques like functional near-infrared spectroscopy warrant consideration.

### Conclusion

We show that people have an expectation of the shape and orientation of an object’s trajectory that is based on natural motion under the laws of gravity. This expectation facilitates manual interceptions by enabling faster movements that fall along the final trajectory. Our results corroborate past findings that gravitational acceleration (*g*) and direction (downward) enhance performance across perceptual, gaze, and interception tasks. They also highlight the importance of nonlinear trajectory shapes in modulating expectations, and more specifically, the important contribution of trajectory shape for the interception of projectiles and bouncing objects.

## Acknowledgements

This research was funded by a Natural Sciences and Engineering Research Council of Canada Discovery Grant (04271-2022 RGPIN) to JCC. Ben Chang collected pilot data for a preliminary version of this experiment using a back-projection screen instead of virtual reality. The authors thank Drs. Paul Gribble and Joern Diedrichsen for helpful advice on computing average error ellipses, Kevin Stubbs and Derek Quinlan for technical assistance, and Cassandra Bacher for assistance with ethics board reporting.

