## Supplementary Figures for "Target interception in virtual reality is better for natural versus unnatural trajectory shapes and orientations"

### Supplementary Materials

#### Accuracy

A three-factor ANOVA was performed on the data for accuracy. In addition to a two-way interaction between *Shape* and *Orientation* and a main effect of *Shape* (discussed in the main text), this analysis identified an interaction between *Orientation* and *Visual Field* ($F_{1,23} = 10.4$, $p = .004$, $\eta^2_p = .31$). As shown in Supplementary Figure 1, this interaction is explained by higher accuracy during the upright LVF condition compared to the other three conditions.

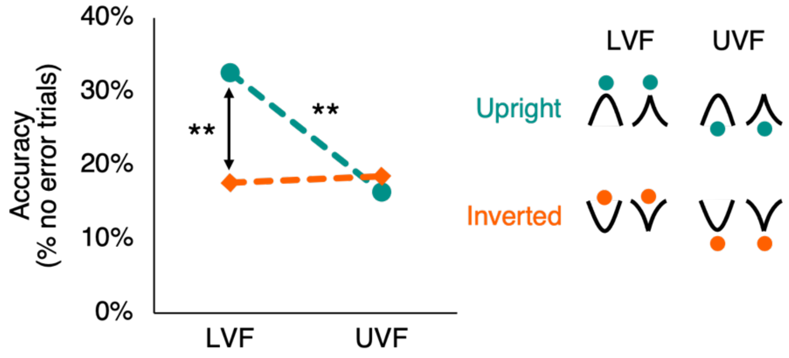


**Supplementary Figure 1. Accuracy across conditions, interaction between *Orientation* and *Visual Field.*** Results here are collapsed across shape. Participants made fewer errors when the trajectory was upright and within the LVF. Accuracy was similar across the other three conditions. The schematic on the right depicts where participants are fixating, represented as the teal and orange circles, relative to the trajectory for each of the four conditions shown. * $p < .05$, ** $p < .01$, *** $p < .001$ after *post-hoc* analysis.

#### Constant Error, y Direction

A three-factor ANOVA was performed on the data for the average y position of the constant error (the average point of interception). As discussed in the main text, this analysis identified a two-way interaction between *Shape* and *Orientation*, and a main effect of *Orientation*. However, this analysis also revealed a three-way interaction between *Shape*, *Orientation*, and *Visual Field* ($F_{1,23} = 14.03$, $p = .001$, $\eta^2_p = .38$) as shown in Supplementary Figure 2a, and a two-way interaction between *Orientation* and *Visual Field* ($F_{1,23} = 30.85$, $p < .001$, $\eta^2_p = .57$) as shown in Supplementary Figure 2b.

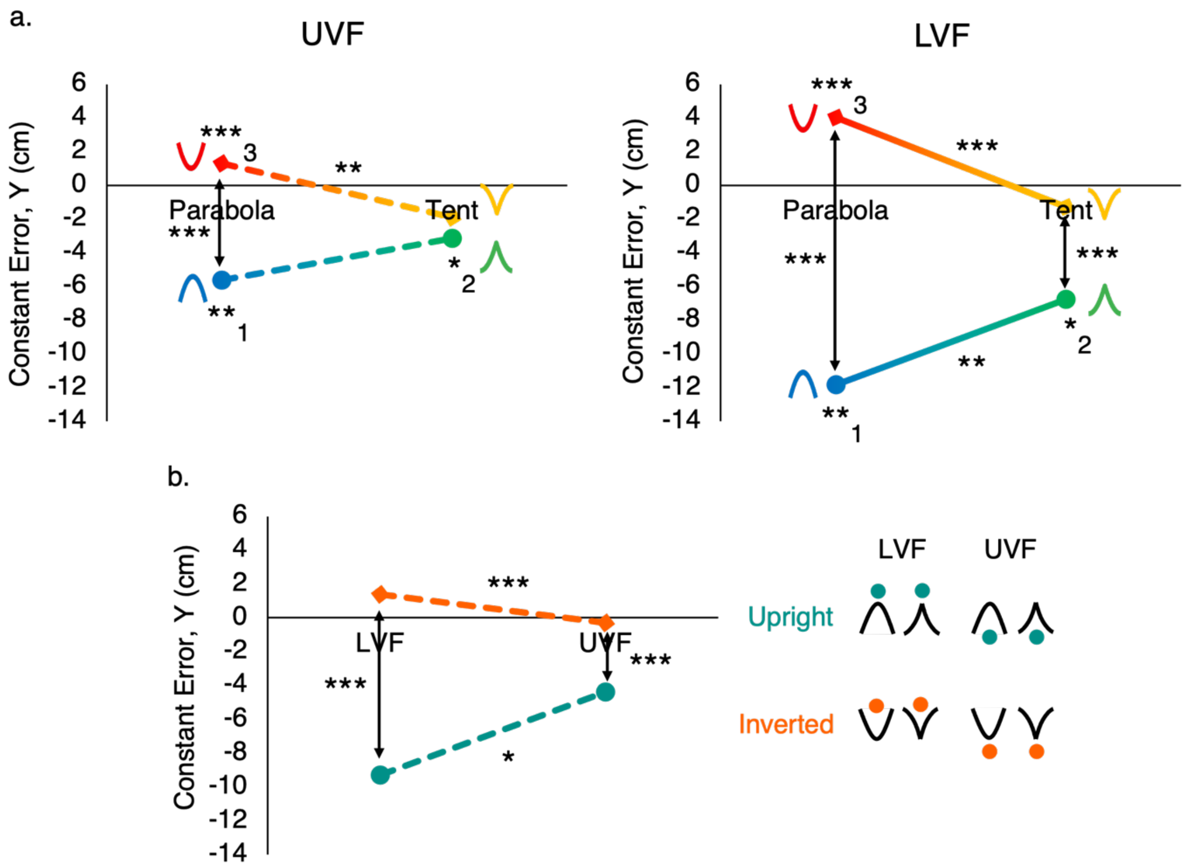


**Supplementary Figure 2. Interactions identified for constant error along the y direction.** * $p < .05$, ** $p < .01$, *** $p < .001$ after *post-hoc* analysis. **(a)** A three-way interaction between *Shape*, *Orientation*, and *Visual Field* was identified. The relationship between *Orientation* and *Shape* is similar regardless of whether the trajectories took place in the LVF or the UVF, however this relationship is more pronounced for UVF trajectories. **(b)** A two-way interaction between *Orientation* and *Visual Field* was identified. Participants generally aimed below the ball for upright trials and above the ball for inverted trials, however they aimed lower below and higher above respectively when the trajectory took place in the LVF compared to the UVF.

The three-way interaction did not qualitatively change the interpretation of our main finding that the interaction between *Shape* and *Orientation* revealed that reaches were generally below the ball for upright trajectories, a relationship that was more pronounced for parabolas. As shown in Supplementary Figure 2a, reaches were on average lower for upright compared to inverted trajectories for trajectories in both the LVF and UVF. This relationship is more pronounced for trajectories in the LVF, wherein reaches for upright trials were considerably lower on average than reaches for upright trials in the LVF. Similarly, reaches for inverted parabola trials were higher for LVF compared to UVF trials. A possible explanation for this finding is that since interceptions in the LVF occurred in a more natural condition, this may have evoked stronger expectations of real-world conditions, such that participants were more likely to aim below the ball to intercept upright targets and aim above the ball to intercept inverted targets, which is a sensible approach in the real world. While the relationship is stronger for the LVF compared to the UVF, the main take aways did not qualitatively change based on this finding. Similarly, Supplementary Figure 2b demonstrates that reaches were generally lower for upright compared to inverted conditions regardless of the visual field.

**Movement Time**

A three-factor ANOVA was conducted on the data for median movement time. As described in the main text, this analysis identified a two-way interaction between *Shape* and *Orientation* as well as a main effect of *Shape*. However, this analysis also identified a two-way interaction between *Orientation* and *Visual Field* ($F_{1,23} = 8.8$, $p = .007$, $\eta^2_p = .28$), as shown in Supplementary Figure 3. This interaction is explained by a longer movement time for upright than inverted trajectories in the UVF. Movement time was similar between the remaining three conditions. Although it is unclear why this result emerged, it does not change our main finding that when faced with unnatural trajectories participants compensated for a slower reaction time with a shorter movement time.

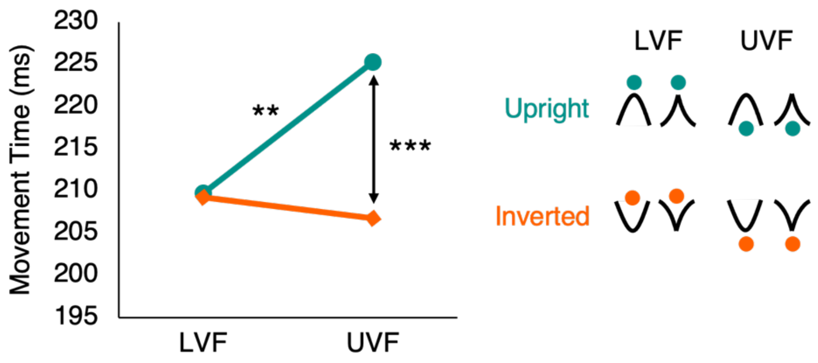


**Supplementary Figure 3. Two-way interaction between *Orientation* and *Visual Field* for movement time.** Results here are collapsed across shape. Movement time was longer when trajectories were upright and in the UVF. Movement time did not differ significantly between the three other conditions. The schematic on the right depicts where participants are fixating relative to the trajectory for each of the four conditions shown. The teal and orange circles represent the fixation locations for upright and inverted trajectories, respectively. * $p < .05$, ** $p < .01$, *** $p < .001$ after *post-hoc* analysis.
